# Genomic analyses of Asiatic Mouflon in Iran provide insights into the domestication and evolution of sheep

**DOI:** 10.1101/2023.10.06.561316

**Authors:** Dong-Feng Wang, Pablo Orozco-terWengel, Meng-Hua Li, Feng-Hua Lv

## Abstract

**Background:** Asiatic mouflon (*O. gmelini*) consists of several subspecies mainly distributed in Armenia, southern Azerbaijan, Cyprus, northern, southern, and western regions of Iran, and eastern and central regions of Turkey nowadays. Genome analyses of Asiatic mouflon in Iran revealed that they could have diverged from the direct ancestor of domestic sheep, and showed genetic introgression into domestic sheep after domestication. However, the impact of the Asiatic mouflon subspecies in Iran on sheep domestication remains unclear.

**Results:** Here, we conducted a comprehensive population genomics analysis of Asiatic mouflons in Iran with 780 whole-genome sequences, 767 whole mitogenomes, and 239 Y chromosomes. Whole-genome sequence analyses revealed two subpopulations of the Asiatic mouflons in Iran: *O. gmelini*_2 limited in Kabudan and *O. gmelini*_1 over a wide geographic area. Phylogenetic analyses of the Asiatic mouflons in Iran based on uniparental variants revealed a monophyletic lineage with the mitochondrial haplogroups C+E, and clustered into a monophyletic with Y-chromosomal lineage HY2 of sheep. Additionally, introgression tests detected significant signals of genetic introgression from *O. gmelini*_2 to four sheep populations (e.g., Garut, Garole, Bangladeshi, and Sumatra) in South and Southeast Asia. In the four sheep populations, selective tests and introgression signals revealed that the wild introgression could have contributed to their small body size and local adaptation to the hot and humid environments in the Indian Peninsula.

**Conclusions:** Our results suggested that the maternal haplogroups C+E and paternal lineage HY2 could have originated from the Asiatic mouflon in Iran. Also, our findings provide new insights into sheep domestication and subsequent introgressions from wild ancestors to domestic populations.

## Background

Archeozoological and genetic evidence suggested that Asiatic mouflon (*O. gmelini*), the wild ancestor of sheep, have underwent a period of multigenerational human management in southeastern Anatolia by ∼10,500 years B.P. [1–4]. Following domestication, sheep diffused throughout Eurasia with human migrations and developed into different populations under natural and artificial selections [5–8]. Asiatic mouflon ranges over a wide geographic area in Armenia, southern Azerbaijan, Cyprus, northern, southern, and western regions of Iran, and eastern and central regions of Turkey. It has been classified into five subspecies based on morphological differences and geographic distributions [9, 10], such as Armenian mouflon (*O. gmelini gmelini*) [11], Anatolian mouflon (*O. gmelini anatolica*) [11], Isfahan mouflon (*O. gmelini isphahanica*), Laristan mouflon (*O. gmelini laristanica*), and Cyprus mouflon (*O. gmelini ophion*) [11]. Due to the complex intraspecies taxonomy of Asiatic mouflon, the genetic contributions of various subspecies to the domestication of sheep remains unclear [3, 12–14].

Previous genetic studies revealed controversial ancestral origins of domestic sheep. Hiendleder et al. (2002) proposed that sheep were domesticated from two different subspecies in Turkey and western Iran, corresponding to mtDNA lineages A and B of domestic sheep, respectively [13]. Through genetic and phylogeographic analyses of partial mtDNA sequences of modern and ancient samples in wild and domestic sheep, Demirci et al. (2013) suggested that domestic sheep could have originated from two distinct ancestral populations of mtDNA lineages A+B+D and C+E of domestic sheep, respectively [3]. Based on genetic, archeozoological, and geographic evidence of 212 Asiatic mouflon samples covering most of their distribution area, a recent study suggested that the maternal origin of sheep could be traced to a subspecies of *O. gmelini gmelini* in eastern Anatolia and central Zagros [4].

Despite the intensive focus on material inheritance, Y-chromosome and whole-genome analyses have received less attention. Recently, Y-chromosome and whole-genome analyses of Asiatic mouflon in Iran revealed that they could have diverged from the direct ancestor of domestic sheep, and showed genetic introgression into domestic sheep after domestication [8, 14, 15]. Also, these studies have demonstrated that genomes of Asiatic mouflon in Iran are crucial for the development of morphological and adaptive characteristics in domestic sheep [8, 15, 16]. Nevertheless, the impact and extent of genetic introgression involving the Asiatic mouflon subspecies in Iran (e.g., *O. gmelini gmelini*, *O. gmelini isphahanica*, and *O. gmelini laristanica*) and their hybrid populations (*O. gmelini gmelini* × *O. vignei*) on the genomes of domestic sheep remains unclear.

Here, we comprehensively analyzed whole genome sequence, complete and partial mtDNA sequences, and Y-chromosome genetic variants in domestic sheep and Asiatic mouflon, covering a large distribution area in Asia, Europe, Africa, and America. We aimed to elucidate the contributions of various Asiatic mouflon subspecies in Iran to the domestication of sheep. Additionally, we investigated the impact of wild introgression of the Asiatic mouflon subspecies in Iran on the genomes of domestic sheep populations in South and Southeast Asia, which have adapted to the hot and humid environments near the Indian Ocean.

## Methods

### Datasets

Whole-genome sequences (WGS) of 780 individuals representing 158 domestic sheep populations and three wild sheep species (32 Asiatic mouflon (*Ovis gmelini*) in Iran, 9 urial (*Ovis vignei*), and 1 bighorn sheep (*Ovis canadensis*)) were obtained from publicly available data [See Additional file 1, Table S1]. Besides these, we retrieved 161 whole mitogenomes from the NCBI [See Additional file 1, Table S2]. Also, we included 333 Cytochrome b (*Cytb*) sequences of Asiatic mouflon in the integrated analysis [See Additional file 1, Table S3].

### Variant calling

We implemented quality-based trimming for WGS data with Trimmomatic v0.39 [17] and mapped high-quality reads to the reference genome *Oar_rambouillet_v1.0_CM022046.1*. (https://www.ncbi.nlm.nih.gov/assembly/GCF_002742125.1/; https://www.ncbi.nlm.nih.gov/nuccore/CM022046.1/, last accessed May 12, 2020) using the Burrows–Wheeler aligner (BWA mem) v.0.7.8 [18] with the default parameters. Then, we removed duplicates with the *MarkDuplicates* module in GATK v 4.1.2.0 [19] and called SNPs following the previous procedures [15]. We used the *HaplotypeCaller* module in GATK v 4.1.2.0 to generate the GVCF file of each sample and merged the raw GVCF file of each individual with the *CombineGVCFs* module. Afterward, we genotyped SNPs using the *GenotypeGVCFs* module and filtered SNPs using the *VariantFiltering* module of GATK with the parameters “QUAL < 30.0 || QD < 2.0 || MQ < 40.0 || FS > 60.0 || SOR > 3.0 || MQRankSum < - 12.5 || ReadPosRankSum < -8.0”. We obtained a total of 121,252,917 high-quality autosomal SNPs and 203,938 Y-chromosomal SNPs for the following analyses.

### Mitochondrial DNA assembly

We assessed the mitogenome depth of each sample using the SAMtools v.1.9 [20] and kept only the samples with depth > 100 in the below analyses. We assembled the complete mitogenomes using the following protocols. We first converted BAM files to the fastq format. Then, we generated the mitogenome consensus sequence from the fastq files using MIA v1.0 (https://github.com/mpieva/mapping-iterative-assembler).

In addition to the 605 mitogenomes assembled here, we also included 161 whole mitogenome sequences retrieved from the NCBI GenBank in the analysis, with a final mitogenome dataset of 767 sequences from 727 domestic sheep, 31 Iranian Asiatic mouflon, and 9 urial. All sequences were aligned using MAFFT v7.515 [21] and trimmed with trimAl v1.4 [22].

### Y-chromosomal SNPs

To obtain single-copy Y-chromosomal SNPs, raw SNPs in the 780 samples were processed as follows. Firstly, we identified ram samples from the whole samples using PLINK v1.90b6.26 [23] with the “--check-sex” option and kept only these ram samples in the following analyses. We then excluded SNPs in the regions covering the pseudoautosomal regions (PARs) and the highly repetitive sequences (around 1.06 Mb) [14]. Afterwards, we filtered SNPs with the mean sequencing depth ranging from 0.5 × to 2 × of the expected depth (half of the genome-wide depth) [14]. Further, we removed SNPs that were heterozygous in at least one of the male samples. Finally, we retained 4,096 Y-chromosome SNPs for the haplotype and phylogenetic analyses.

### Population structure analysis

To reveal the genetic differentiation within the Asiatic mouflon in Iran, we examined the population genetic substructure for the total 779 wild and domestic samples using a set of autosomal SNPs filtered as follows. First, we filtered the total 25,270,188 high-quality SNPs with missing rates < 0.1 and minor allele frequencies > 0.05 (MAF > 0.05) using PLINK v1.90b6.26 [23]. Then, we pruned the remaining SNPs with the options “--indep-pairwise 50 10 0.1” of PLINK v1.90b6.26 [23]. Afterwards, 1,566,300 SNPs were retained for the population genetic structure analysis.

We computed the identity by state among samples using PLINK v1.90b6.26 [23] with option “distance square 1-ibs”. Next, we generated a neighbor-joining (NJ) tree using the function “bionj” in the R package *ape* v5.6.2 [24]. The tree was rooted using urial (*O. vignei*) as the outgroup, and its graphical representation was obtained using the R package *ggtree* v3.6.2 [25]. Principal component analyses (PCA) were performed using the “--pca” option in PLINK v1.90b6.26 [23] with the default settings. Furthermore, model-based clustering was performed using the sNMF v1.2 with clusters *K* from 2 to 10 [26].

The *F*_ST_ and nucleotide diversity (π) of autosomes were calculated with vcftools v0.1.17 [27] using 2Mbp windows and 1Mbp steps, and *P* value was calculated with 1000 permutation test. The *F*_ST_ of mitochondria and Y chromosome were calculated with R package hierfstat v0.5.11 [28] with 1000 permutation test. The haplotype diversity of mitochondria and Y chromosome were calculated with pegas v1.1 [29] with 1000 permutation test.

### Phylogenetic analysis

Phylogenetic analyses based on mitogenome and Y-chromosomal SNPs were performed using the maximum likelihood (ML) algorithm of iqtree2 v2.0.7 [30]. Mitogenome sequences were aligned using mafft v7.515 [21] with the default setting, and multiple sequence alignment (MSA) was trimmed to remove ambiguously aligned regions using trimAl v1.4 [22] with parameters “-automated1”. To keep high-quality SNPs, we further removed all the sites with gaps and ambiguous nucleotide bases and only kept parsimony informative sites using *ClipKIT* [31]. We tested the potential models of DNA sequence evolution using the option “-m” test in iqtree2 v2.0.7 [30] and found the generalized time reversible model (GTR + G) [32] was the best fitting substitution model for mitogenome. We constructed a maximum likelihood (ML) tree with the site model GTR + G with 1000 bootstrap replicates using iqtree2 v2.0.7 [30]. The resulting tree was rooted with bighorn as the outgroup and visualized using the FigTree v1.4.4 (http://tree.bio.ed.ac.uk/software/figtree/). Mitogenome haplotypes were inferred with the function “haplotype” in the R package *pegas* v1.1 [29]. We also constructed the parsimony mitogenome haplotype network using POPART [33]. We transformed the genotypes of Y-chromosomal 4,096 SNPs into the fasta format using vcf2phylip.py [34]. Phylogenetic trees were built and visualized following the same pipelines and using the same programs and substitution model as above.

### Demographics and divergence time inference

The population genetic structure analyses above identified two subpopulations (*O. gmelini*_1 and *O. gmelini*_2) in the Asiatic mouflons in Iran. We inferred the dynamic changes in historical and recent effective population size (*N*_e_) of the two subpopulations using SMC++ v1.15.2 algorithms [35]. We selected 7 - 10 high-depth individuals in each of the six geographically differentiated genetic groups of domestic sheep (Central-and-East Asian, South-and-Southeast Asian, the Middle Eastern, African, European, and American populations) [15] and the two Asiatic mouflon subpopulations. We estimated the *N*_e_ during the past 1,000 to 100,000 years for each group using SMC++ v1.15.2 [35] under the default setting.

For the SMC++ analysis, the reference genome’s low-complexity regions were masked using snpable (http://lh3lh3.users.sourceforge.net/snpable.shtml, last accessed October 20, 2022) using in-house scripts. We set the mutation rate and generation time to be 1.00×10^-8^ per nucleotide site per generation [36] and 3 years [37, 38], respectively. To infer the divergence times between *O. gmelini*_1 and *O. gmelini*_2 as well as between the Asian mouflon and domestic sheep, we employed the split option of SMC++ with the same parameter settings as above.

Additionally, we inferred the demographic history of domestic sheep and Asiatic mouflon based on mitogenomes and Y-chromosome data using Beast v2.7.3 [39] and the Coalescent Bayesian Skyline plot [40]. Firstly, we used a coalescent model to estimate the mutation rates for mitogenome and Y chromosome chromosomes. The analysis was performed using the GTR + GAMMA (4) + I site model and the strict clock model. The most recent common ancestor (MRCA) of Asiatic mouflon and domestic sheep was set to be 0.653 million years [41]. The chain length was set to be 40,000,000 with a burn-in of 2,000,000. The results were analyzed using Tracer (http://beast.community/tracer), assuming mutation rates estimates of 2.10 × 10^-8^ and 7.70 × 10^-9^ substitutions per site per year for the mitochondrial and Y chromosome SNPs, respectively.

Subsequently, we utilized the mutation rates estimated above to infer the demographic history of maternal and paternal lineages in domestic sheep and Asiatic mouflon. We used the same GTR + GAMMA (4) + I site model as above, with the clock model set as strict and the mutation rates of 2.10 × 10^-8^ and 7.70 × 10^-9^ per site per year for mitochondria and Y-chromosome, respectively. The prior model remained the Coalescent Bayesian Skyline, with the chain length of 40,000,000, a burn-in of 2,000,000, and samples drawn every 4,000 steps. The obtained results were further analyzed using Tracer, and each analysis yielded over 200 effective samples for all relevant parameters.

### Introgression analysis

We conducted introgression analysis to clarify the genetic contributions of the two Asiatic mouflon subpopulations to South-and-Southeast Asian (SSA) sheep. We tested introgression using a combination method of *D* and *f*_dM_ statistics. Specifically, we calculated *D* statistics [42] with a four taxon model (((P1, P2), P3), P4) using Dsuite v0.5r50 [43]. In this analysis, Menz sheep (MEN) was selected as the reference population (P1), while SSA populations were the target population (P2), *O. gemlini*_1 or *O. gemlini*_2 was the donor (P3), and *O. canadensis* was the outgroup (P4). A *P* value < 0.05 from a standard block-jackknife procedure was considered as evidence of gene flow from P3 to P2.

We further implemented sliding window statistics analysis using the Dsuite v0.5r50 [43] to quantify the introgressions. We calculated *f*_dM_ values using a sliding window of 50 SNPs with a step of 25 SNPs across the genomes with the defined trios (MEN, SSA, *O. gmelini*_2), where SSA represents the four South-and-Southeast Asian sheep populations suggested to be the introgressed populations by the *D*-statistics and *f*_4_-ratio tests. We calculated the Z-transformed *f*_dm_ values, and the windows with *P* < 0.05 and *D* > 0 was defined as the potential introgressed genomic regions. We evaluated the statistical significance using one-tailed *Z*-test in R package BSDA v1.2.1 [44].

### Selective sweep tests

To assess the potential roles of genomic introgression from Iranian Asiatic mouflons in Southern and Southeastern Asian (SSA) sheep populations, we conducted a PBS analysis [45] to identify selective signatures in four introgressed populations (i.e., Garut (GUR), Garole (GAR), Bangladeshi (BGE), and Sumatra (SUM)). We pooled these four populations (GUR, BGE, SUM, and GAR) as an introgressed group, and used the MEN and *O. gmelini*_2 as the reference populations. We calculated the PBS values using PBScan v2020.03.16 with parameters “-win 50 -step 25” described in Hämälä and Savolainen (2019) [46] and estimated *P* values by 2000 permutation cycles for Monte Carlo testing. and detected selective regions with top 1% PBS values. Subsequently, we identified the overlaps between the selective and the introgressed regions using BEDtools v2.30.0 [47] and annotated functional genes in the overlapping genomic regions.

### Gene annotation and function analyses

We annotated genes in the introgressed regions using the sheep reference assembly *Oar_rambouillet_v1.0* by BEDtools v2.30.0 [47]. We also used the sheepQTLdb (https://www.animalgenome.org/cgi-bin/QTLdb/OA/index) [48] to detect QTLs associated with the introgressed regions. We examined the LD blocks by LDBlockShow v1.40 [49]. The LD decay was calculated with PopLDdecay v3.42 [50]. *d*_xy_ and *F*_ST_ for the introgressed region is calculated with Pixy 1.2.7.beta1 [51] with a 20kb sliding window with 2kb steps. The haplotype heatmap of regions near the introgression region was plotted with R package ComplexHeatmap v2.14.0 [52].

## Results

### Genetic variants, mitogenome assembly, and Y-chromosome specific SNPs

A total of 121,252,917 SNPs in the 780 wild and domestic sheep samples were obtained for downstream analyses. We obtained an average depth of 923.33 × (101.92 × - 8,324.72 ×) for the mitogenome assemblies after aligning all mitochondrial reads from the 606 samples to the reference mitochondrial genome NC_001941.1 [See Additional file 1, Table S2]. After quality control for the SNP identification on the Y-chromosome, we identified 4,096 Y-specific SNPs in a cohort of 239 rams with an average read depth of 8.26 × (∼4.25 × - 16.99 ×) for SNP [See Additional file 1, Table S4].

### Population genetic structure of the Asiatic mouflons in Iran

We observed two subpopulations in the 32 Asiatic mouflons in Iran (Fig. 1) with autosome SNPs. The larger subpopulation (*O. gmelini*_1) comprised 25 samples across eastern and southern Iran, while the smaller subpopulation (*O. gmelini*_2) consisted of only 7 samples limited in northwestern Iran (Fig. 1a) and [See Additional file 1, Table S2]. Low kinship values within two subpopulations eliminate inbreeding caused by sampling [See Additional file 1, Table S5]. We also observed significant genetic divergence and different genetic diversity patterns [See Additional file 1, Table S6] and [See Additional file 1, Table S7]. The result makes sure that the divergence is not the artefact of inbreeding.

**Figure 1.**
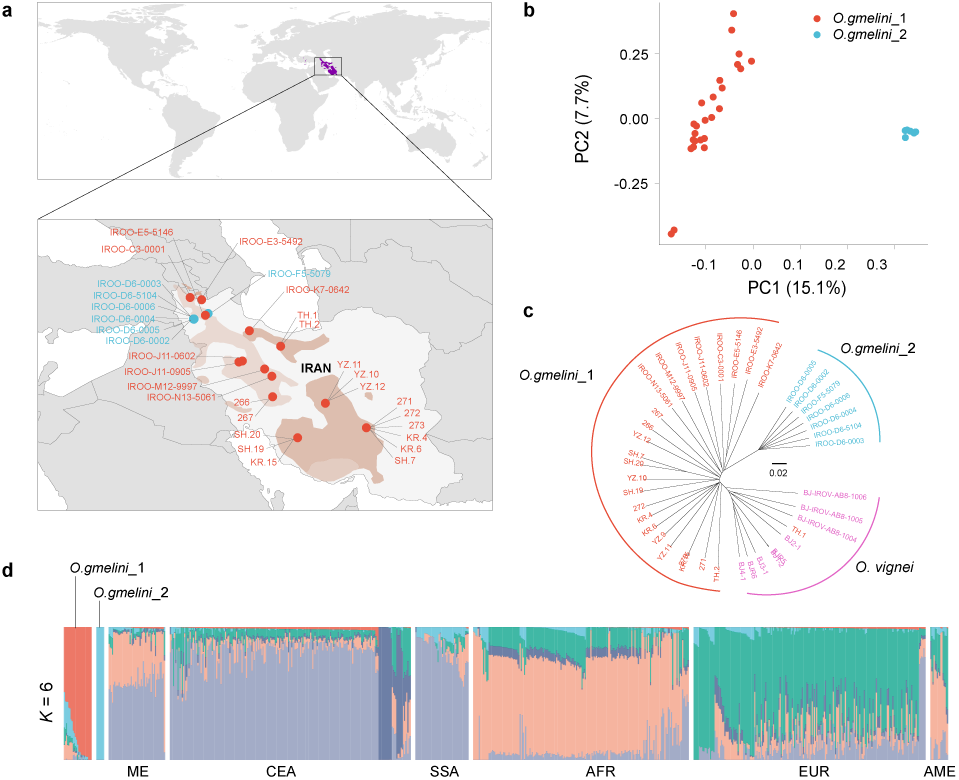
Population genetics structure of Asiatic mouflons in Iran (*O. gmelini*). **a** Geographic distribution of 32 Asiatic mouflons. Purple and blue dots indicate the location of *O. gmelini*_1 and *O. gmelini_2* subpopulations; **b** Principal components 1 and 2 for the 32 Asiatic mouflon in Iran; **c** Neighbor-joining (NJ) trees of the 32 Asiatic mouflons in Iran based on whole-genome SNPs using the nine urial samples as the outgroup; **d** Population genetic structure of the Asiatic mouflons in Iran and domestic sheep inferred from the program the sNMF v1.2 (*k* = 6) using whole-genome SNPs.

Phylogenetic analyses of the total 780 wild and domestic sheep samples identified the same two subpopulations of Asiatic mouflon (*O. gmelini*_1 and *O. gmelini*_2) and six genetic clusters of domestic sheep, which were congruent with their geographic distribution [See Additional file 2, Figure S1] and [See Additional file 2, Figure S2]. Admixture analysis showed inter- and intra-species genetic divergence similar to the phylogenetic tree (Fig. 1d).

### Phylogenetic and haplotype analysis based on uniparental variants

We observed 593 mitochondrial haplotypes from the 767 mitogenomes of wild and domestic sheep (Fig. 2a) and [See Additional file 1, Table S2]. The phylogenetic tree inferred from the mitochondrial haplotypes revealed three monophyletic clades (clades I, II, and III) (Fig. 2b). Clade I comprised of haplogroups A, B, and D of domestic sheep. Clade II consisted of the domestic sheep’s haplogroups C and E and the Asiatic mouflon samples, while the remaining Asiatic mouflon samples clustered with the urial in clade III (Fig. 2b).

**Figure 2.**
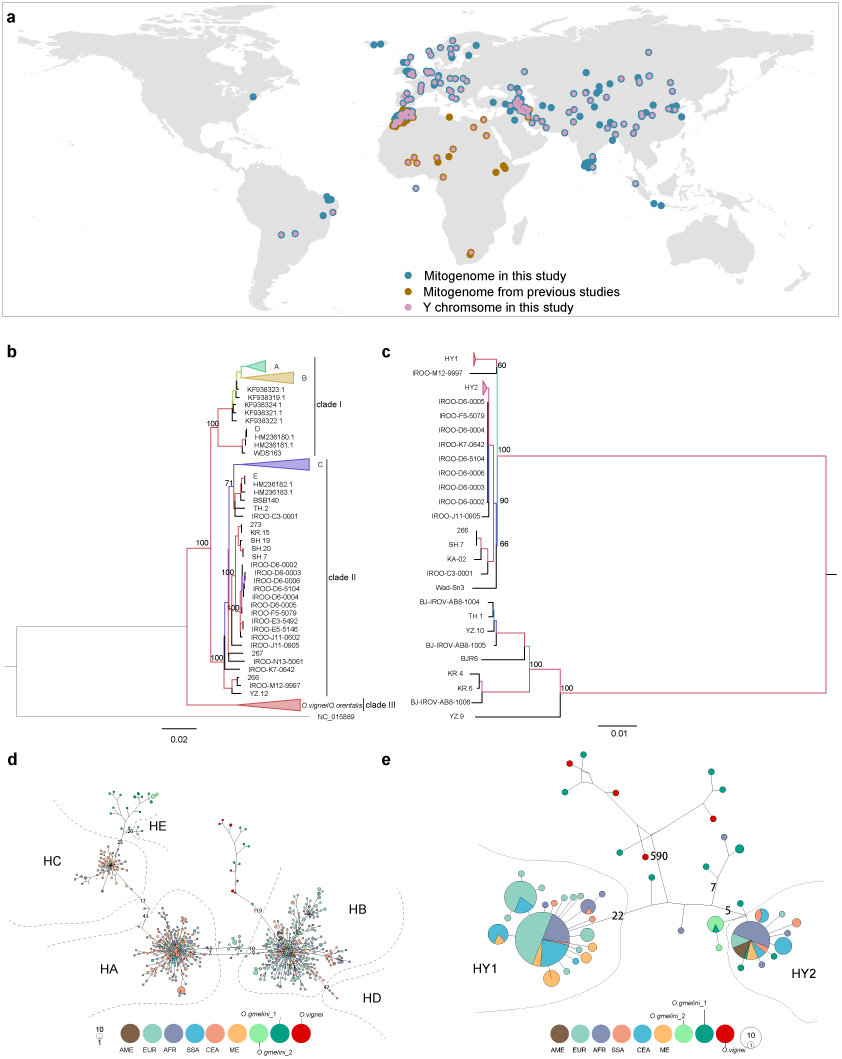
Phylogeny of Asiatic mouflon, urial, and domestic sheep inferred from mitogenomes and Y-chromosomal genetic variants. **a** Geographical distribution of the samples in this and previous studies; **b** Phylogeny of Asiatic mouflon, urial, and domestic sheep inferred from 767 mitogenomes; **c** Phylogeny of Asiatic mouflon, urial, and domestic sheep inferred from 239 Y-chromosomal SNPs; **d** Network of mitogenome haplotypes; **e** Network of Y-chromosome haplotypes. Bootstrapping values above 50% were labeled on the phylogenetic tree. Bighorn sheep were used as the outgroup in phylogenetic analyses based on mitogenome and Y-chromosome genetic variants.

For Y-chromosomal SNPs, the lineage HY2 of domestic sheep was closely related to Asiatic mouflon in the haplogroups C and E but diverged from the main lineage HY1 (Fig. 2c) and [See Additional file 1, Table S8]. However, phylogenetic and network analysis revealed no shared mitochondrial haplotypes and only one shared Y-haplotypes of HY2 between domestic sheep and Asiatic mouflon in Iran (Fig. 2d and e). We observed significant genetic divergence (*O. gmelini*_1: *F*_ST_ = 0.11, *P* < 0.05; *O. gmelini*_2: *F*_ST_ = 0.21, *P* < 0.05) between Asiatic mouflons and domestic sheep in whole genome levels [See Additional file 1, Table S6]. The observation implied that the Asiatic mouflon in Iran might not be the direct ancestor of domestic sheep. Instead, the relatively close phylogenetic relationships between the Asiatic mouflons in Iran and domestic sheep suggested by haplogroups C, E, and Y-chromosomal haplogroupY2 implied their possible gene flows.

### Effective population size and divergence time

Inference of demographic history showed the Asiatic mouflons and domestic sheep shared a similar historical pattern 10,000 years ago (Fig. 3a), confirming the Asiatic mouflon as the wild ancestor of sheep [53]. SMC++ analysis revealed a continual decline of *N*_e_ in domestic sheep and Asiatic mouflon during ∼8,000–10,000 years ago. Nevertheless, the two Asiatic mouflon populations in Iran showed slower declines than domestic sheep (Fig. 3a), which probably reflected the bottleneck effect of domestication. A sharp decline of *N*_e_ was detected ∼1,000-5,000 years ago in the *O. gmelini*_2, demonstrating a pattern different from that for the *O. gmelini*_1 (Fig. 3a). We observed similar estimates of divergence time between *O. gmelini*_1 and the six geographically differentiated genetic groups of domestic sheep during ∼9,000-15,000 years ago (Fig. 3b) and [See Additional file 1, Table 9], which was congruent with the domestication time of sheep [2]. Additionally, we found the splitting time between the two Asiatic mouflon populations in Iran around 5,000 years ago (Fig. 3b) and [See Additional file 1, Table 9], indicating a recent genetic divergence of Asiatic mouflons in Iran (Fig. 1). For the divergence pattern inferred by WGS of the Asian mouflons, the subpopulation *O. gmelini*_2 clustered with partial individuals of the subpopulation *O. gmelini*_1 in the mitogenome and Y-chromosomal phylogenetic trees (Fig. 1 and 3). The discordant phylogenetic patterns of Asiatic mouflons in Iran inferred by uniparental and autosomal genetic variants could be explained by the recent hybridization events [54].

**Figure 3.**
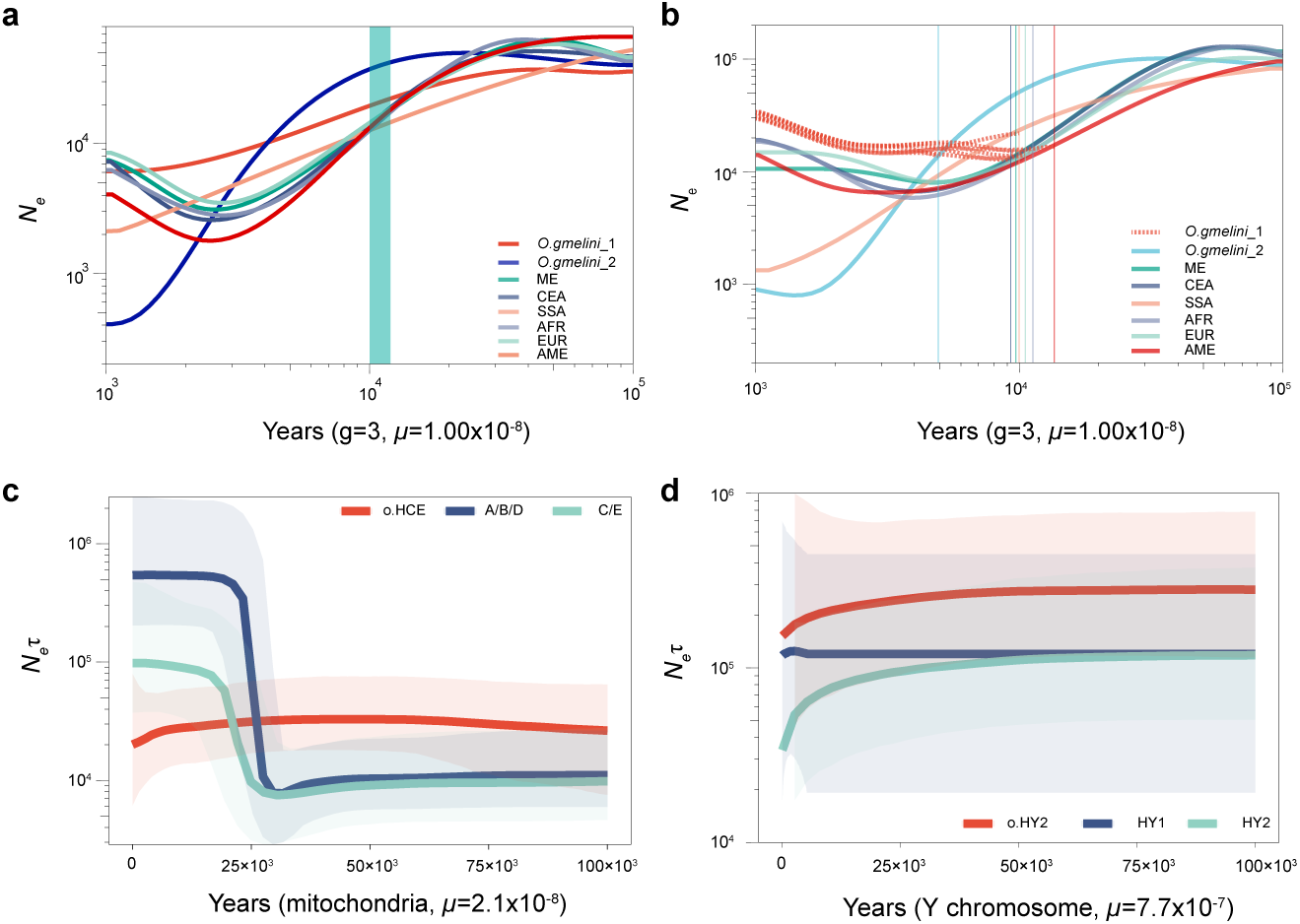
Demographic history of Asiatic mouflon in Iran and domestic sheep. **a** Effective population sizes (*N*_e_) across six domestic sheep populations and the two subpopulations of Asiatic mouflon in Iran inferred from SMC++; **b** Inference of split times between *O. gmelini*_1 and the other wild and domestic sheep populations. The vertical lines represent the split time between each pair of populations; **c** Bayesian skyline plot (BSP) reconstruction of historical change in effective population size (*N*_e_) of domestic sheep of maternal haplogroups HA+B+D, and HC+E, and the Asiatic mouflons in Iran; **d** Bayesian skyline plot (BSP) reconstruction of historical changes in *N*_e_of paternal lineages HY1 and HY2 of domestic sheep and the Asiatic mouflon in Iran. The middle line represents median estimates of *N*_e_τ, and the shaded area represents 95% highest posterior density (HPD) intervals.

We conducted Bayesian skyline analyses to estimate *N*_e_ based on mitogenomes and Y-chromosomes. Based on the estimated TMRCA (time to the most recent common ancestor) of Asiatic mouflons and domestic sheep [41], three lineages (lineage o.HCE: haplogroup of Asiatic mouflons in Iran and partial domestic sheep of haplogroups C and E, lineage A+B+D: haplogroups A, B, and D, and lineage C+E: haplogroups C and E) of mitogenome showed different patterns. Lineage A+B+D showed a steep increase in *N*_e_ at approximately ∼25 k years ago. For lineage C+E, it also increased at approximately ∼25 k years ago, while lineage o.HCE demonstrated relatively fewer changes since approximately ∼100 k years ago (Fig. 3c). Also, we observed different patterns in the three Y-chromosomal lineages (lineage o.HY2: haplogroups of Asiatic mouflons in Iran and partial domestic sheep of HY2, lineage HY1, and lineage HY2). Lineages o.HY2 and HY2 displayed slow decline in *N*_e_ at approximately ∼25 k years ago, while lineage HY1 changed sightly since approximately ∼100 k years ago (Fig. 3d). Thus, uniparental genetic markers showed similar demographic patterns between o.HY2 and HY2 as well as between HCE and C/E, but different patterns between the dominant linages A+B+D and HY1 (Fig. 2 and 3). The results suggested that the Asiatic mouflons in Iran might not be the direct ancestor of domestic sheep but be involved in the genetic introgression.

### Phylogeography of Asiatic mouflon

We built a *Cyt-b* dataset consisting of 247 Asiatic mouflons and 8 European mouflons. The samples of Asiatic mouflon covered the current distribution of all the subspecies (Fig. 4a) and [See Additional file 1, Table S3]. To clarify phylogenetic relationships between different Asiatic mouflon subspecies and domestic sheep, we performed phylogenetic analysis of *Cyt-b* sequences of Asiatic mouflons and domestic sheep using argali as the outgroup. In the phylogenetic tree built from *Cyt-b*, Asiatic mouflon was assigned to 7 major clades supported by high bootstrap values (94∼100) (Fig. 4b). The topology confirmed the five mitogenome clades of domestic sheep. Haplogroups C+E clustered into monophyletic clade, which diverged from the haplogroups A+B+D (Fig. 4b). It was noteworthy that most of the hybrid samples inferred above were integrated with *O. gmelini* or *O. vignei* samples, and only around 10% of samples clustered with subspecies such as *O.g.ophion* and *O.g.anatolica* (Fig. 4b) and [See Additional file 1, Table S10]. Clade 3 comprised haplogroups C+E of domestic sheep and the Asiatic mouflons in Iran with high frequencies of shared haplotypes between them (Fig. 4c) and [See Additional file 1, Table S10]. Haplogroups A+B+D of domestic sheep and the Asiatic mouflons from Iran and Turkey fell within clade 6, but no common haplotypes were detected between domestic and wild sheep (Fig. 4c) and [See Additional file 1, Table S10]. The results were in accordance with the mitogenome analyses and implied that partial haplotypes C+E in domestic sheep could have originated from the Asiatic mouflons in Iran by introgression.

**Figure 4.**
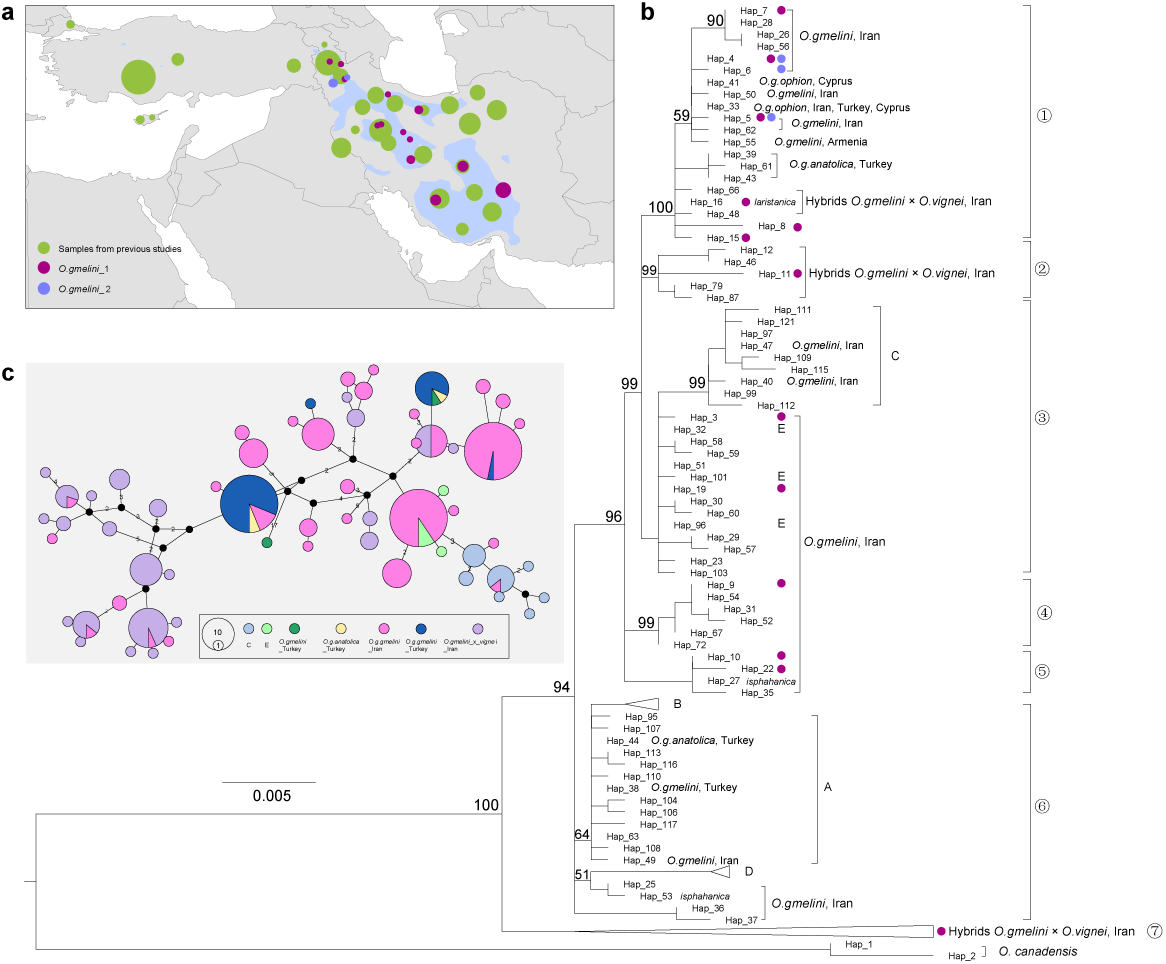
Geographic distribution of Asiatic mouflon and the phylogenetic relationship among Asiatic mouflon subspecies and domestic sheep. **a** Geographical distribution of Asiatic mouflon and samples information of 32 Asiatic mouflon in Iran and 301domestic sheep samples; **b** Phylogenetic tree inferred based on complete *Cyt-b* sequences using the maximum likelihood algorithms. Bootstrap values above 50% were labeled on the phylogenetic tree. **c** Network of Asiatic mouflon in Iran and Turkey and domestic sheep lineage of C and E built from complete *Cyt-b* sequences.

### Genetic introgression

To investigate the impact of introgression from the Asiatic mouflons in Iran on domestic sheep, we implemented introgression tests between Iranian Asiatic mouflon and SSA by *D* and *f*_4_-ratio statistics [55, 56]. We first conducted an admixture analysis between Asiatic mouflons and the SSA populations. We identified three genetic clusters with the smallest cross validation error (*K* = 3) (*O. gmelini*_1, *O. gmelini*_2, and SSA) and detected genetic components of Asiatic mouflons in Iran in some domestic populations (Fig. 5a) and [See Additional file 2, Figure S3]. Further, we employed the ABBA model (((P1, P2), P3), P4) to assess introgression between *O. gmelini*_1 or *O. gmelini*_2 (P3) and SSA sheep populations (P2). We used Menz sheep (MEN) from Africa as P1, which did not show genetic introgression from Asiatic mouflon and SSA populations [8, 15], and bighorn sheep as the outgroup P4, which inhabit North America and have no gene flows with SSA sheep populations. Of the two Asiatic mouflon populations, *O. gmelini*_1 or *O. gmelini*_2 served as P3 and 18 SSA populations as P2. We calculated the estimates of Patterson’s *D*-statistic and *f*_4_-ratio for all the 18 SSA sheep populations, and determined introgression based on the following criteria: *D* > 0, *P*-value < 0.05, and *f*_4_-ratio > 0. Only four sheep populations (Garut, Garole, Bangladeshi, and Sumatra sheep) showed genetic introgression from *O. gmelini_*2 (Fig. 5b) and [See Additional file 1, Table S11]. All the four sheep populations are distributed in the regions near to the Indian Ocean.

**Figure 5.**
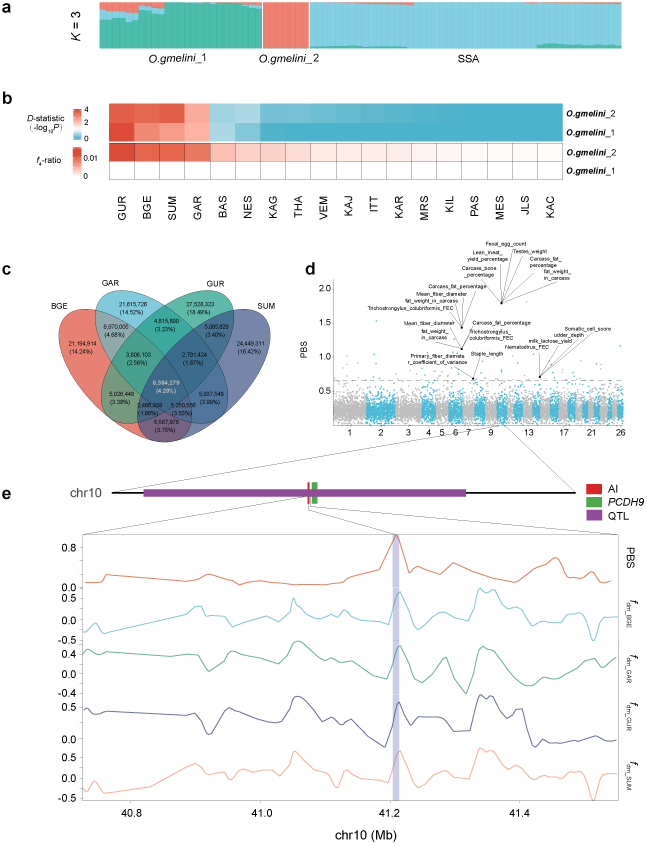
Genetic introgression of Asiatic mouflon in SSA domestic populations. **a** Admixture proportions of Asiatic mouflon in Iran and SSA domestic populations inferred from the program the sNMF v1.2 (*K* = 3) using whole-genome SNPs. **b** Heatmap of *D*-statistics and *f*_4_-ratio calculated by Dsuite v0.5 applied to data including *O. gmelini*_1, *O. gmelini*_2, and SSA domestic populations; **c**, Venn diagram for the shared introgression regions among four domestic South and Southeast populations; **d** Manhattan plot showing signals of selection in the four introgressed SSA populations (BGE/GAR/GUR/SUM) detected by the PBS with the dashed line marking 99.9% threshold. The introgression region, QTL, and adjacent gene *PCDH9* were labeled with blue, purple, and green bars, respectively. The sheep QTLs overlapped with the top 1% PBS regions are shown, **e** PBS and *f*_dm_ values around the significant introgressed genomic region on Chr.10.

To further examine the genetic introgression in the four sheep populations, we calculated *f*_dm_ with a 50-SNP window and 25-SNP step across the sheep genome. We identified introgressed regions genomic with top 5% *f*_dm_ and *D* > 0. We obtained a common introgression region in the four populations spanning 6,384,279 bp (Fig. 5c), [See Additional file 1, Table S12] and [See Additional file 1, Table S13]. We then calculated the PBS value of the four populations to identify regions under selection with MEN sheep and *O. gmelini*_1 as the reference populations. We considered the top 1% of PBS regions as being under selection (Fig. 5d) and [See Additional file 1, Table S14]. We overlapped the selected regions with the common introgressed regions and detected 19,614 bp region as the selected introgressed region [See Additional file 1, Table S15]. We further overlapped the introgressed genomic region with sheep QTLs [See Additional file 1, Table S15]. We found that it overlapped with the QTLs for body_weight, carcass_fat, and fat_weight QTLs, which might be associated with the small body size of Garole, Bangladeshi, and Garut sheep [57–59]. Additionally, the Nematodirus_FEC QTL could be related to the phenotypes of Sumatra sheep’s susceptibility to nematode parasitism [55]. The results implied that the adaptive selected introgressed region could have been introgressed from *O. gmelini*_2, and then have undergone the strong selection to facilitate their adaptation to the hot and humid environments near the Indian Ocean.

## Discussion

Traditionally, Asiatic mouflon had been classified into five subspecies using phenotypic traits, such as coat color, horns, and geographic patterns [11]. However, the intraspecies taxonomy has long lacked genetic evidence. Previous phylogenetic analysis based on partial mtDNA divided Asiatic mouflon into several clades [3, 4, 58], different from the taxonomy of five subspecies defined by phenotypic traits and geographic patterns [11]. Here, we identified two genetic clusters of the Asiatic mouflons in Iran based on WGS data: *O. gmelini*_2 limited in Kaboudan, and *O. gmelini*_1 over a wide geographic area in Iran (Fig. 1). We detected the divergence time of two clusters around ∼5,000 years ago (Fig. 3b) and [See Additional file 1, Table S9], which indicated a recent split of them, and was further supported by the inferred phylogenies based on uniparental genetic markers (Fig. 2b and c). All the individuals of *O. gmelini* _2 collected in Kabudan inhabited the overlap geographic region with Armenian Mouflon (*O. g.gmelini*), while individuals of *O. gmelini* _1 were sampled from the distribution areas of several subspecies such as Armenian Mouflon (*O. g.gmelini*), Isfahan mouflon (*O.g.isphahanica*), and Laristan mouflon (*O.g.laristanica*) (Fig. 1a) [10].

These results suggested a significant genetic divergence between Armenian Mouflon and the other subspecies [See Additional file 1, Table S6]. Nevertheless, the absence of phenotypic data and other sample information for the Asian mouflons and a comprehensive sampling throughout Iran limited our in-depth understanding of within-species genetic differentiation in Asiatic mouflon. Also, Asiatic mouflon is under the status of “Near Threatened”, and their population size has declined in the past two decades under competition with domestic sheep, poaching, and habitat deterioration [10, 60]. Therefore, the population genetic structure analyses and delimitation of taxonomic classification based on WGS data are fundamental for conservation decisions and effective conservation management [61, 62].

Genetic variants of uniparental inheritance offer insights into the domestication and evolutionary history of farm animals, such as pigs [63], sheep [5, 14] and goats [64, 65]. In domestic sheep, here we identified five maternal haplogroups (A, B, C, D, and E) in two monophyletic groups (A+B+D and C+E), and two paternal clades (HY1 and HY2) (Fig. 2b and c). We found that haplotypes of most of the Asiatic mouflons in Iran clustered with haplogroups C+E based on mitochondrial marker (24/31) and grouped with haplogroup HY2 based on Y-chromosomal SNPs (12/21) [See Additional file 1, Table S8]. Nevertheless, the Asiatic mouflons in Iran showed higher divergence from the dominant maternal lineages A and B and paternal lineages HY1 (Fig. 2c and d).

Additionally, *Cyt-b* sequences collected from the whole geographic areas of Asiatic mouflon also revealed a closer relationship between Asiatic mouflon in Iran and domestic sheep of maternal haplogroups C+E (Fig. 4). The results supported the hypothesis that mtDNA haplogroups C+E and Y-chromosomal haplogroup HY2 could have originated from Asiatic mouflons in Iran [14]. However, WGS analysis revealed intraspecies substructure in the Asiatic mouflons in Iran, and genetic introgression from the *O. gmelini*_2 showed a larger impact on domestic sheep than *O. gmelini*_1 (Fig. 5). Similar results were observed from Alberto et al., (2018) that IROO individuals were split into subpopulations and had admixture with some domestic sheep [38]. Therefore, Iran’s two Asiatic mouflon populations showed similar phylogenetic patterns of uniparental inheritance but different introgression patterns of WGS, implying multiple introgression events from the Asiatic mouflons to domestic sheep. In summary, our results implied that Asiatic mouflons in Iran might not be the ancestor of domestic sheep [4, 14], but could have contributed to the gene pool of domestic sheep through multiple introgressions. Also, the results implied the complexity of sheep domestication [4, 66].

Recent studies suggested that sheep diffused to South and Southeast Asia via the Middle East [15]. Our results revealed the genetic introgression from Asiatic mouflon into some South-and-Southeast Asian sheep populations, supporting previous findings [8, 15]. Further, we detected genetic introgression in the four sheep populations of Garut, Garole, Bangladeshi, and Sumatra, and were only from *O. gmelini_*2 in Kabudan (Fig. 5b). The introgression event is congruent with the demographic history of *O. gmelini_*2 and South-and-Southeast Asian sheep inferred here and previously: South-and-Southeast Asian sheep descended from Central-and-East Asian populations around 5.85 to 6.01 ka and then admixed from the Middle Eastern sheep 3.93–4.18 ka [15], and *O.gmelini_*2 split from *O.gmelini_*1 around ∼5,000 years ago (Fig. 3b).

In the four sheep populations, we found that common introgression signals from *O. gmelini_*2 covered about 4.29% of the total introgressed regions. In contrast to population-specific introgression, common introgression could be related to a specific trait or adaptation to similar environments, such as *EPAS1* for the adaptation to hypoxia [45, 67]. We further integrated *PBS* analyses and found four positive selective regions in the common introgressed regions, which were related to carcass_fat_percentage, lean_meat_yield_percentage (Fig. 5d). This might have contributed to the small body size phenotype of Garole, Sumatra, and Bangladeshi sheep [58, 68, 69]. We also found a candidate gene *PCDH9* downstream ∼679 kb of the AI region (Fig. 5e), which had been reported to be associated with local adaptation (e.g., hot and arid environments) in sheep and goats [70, 71] and fat deposition [72]. The gene showed a higher linkage with the AI region (Fig. 6a and b). The AI region was confirmed by *d*_XY_ and *F*_ST_ of Menz sheep (MEN) and introgressed breeds (BGE, GAR, GUR, and SUM) from SSA (Fig. 6c). We observed similar haplotype patterns among introgressed breeds and *O.gmelini*_2 in the AI region contrasting with MEN.

**Figure 6.**
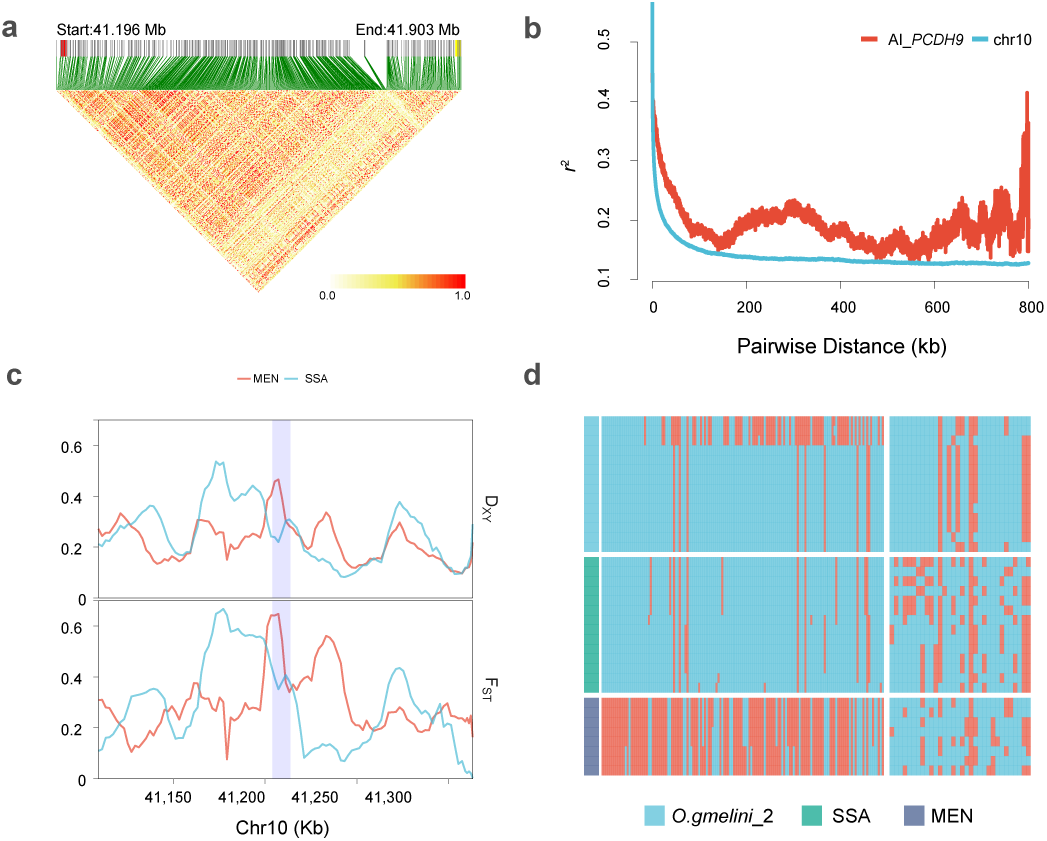
Candidate gene *PCDH9* associated with the significant introgression signal on Chr. 10. **a** Linkage disequilibrium (LD) blocks in the significant introgression region and gene *PCDH9* (chr10:41,104,271-43,073,231). The red and yellow bars represent the location of the significant introgression region and gene *PCDH9*; **b** Pattern of LD decay in the introgressed populations. The black line is calculated from the whole 10 chromosomes, the red line is calculated from the beginning of the significant region to the next 800 kb (chr10:41,204,271-42,004,271); **c** *D*_xy_ and *F*_ST_ around the significant introgression region on Chr. 10 (chr10:41,104,271-41,313,804), calculated by pixy v1.2.7.beta1; **d** Haplotype pattern of the significant introgression region on Chr. 10 (chr10:41,204,271-41,213,804) (left) and exons of *PCDH9* (right).

In contrast, no similar haplotype pattern was found in the exon of *PCDH9* (Fig. 6d). These results implied that the AI region may regulate the function of *PCDH9*. Also, early studies showed adipogenesis, ECM remodeling, inflammation, and lipid droplet dynamics in mature adipose tissue played essential roles in response to climate changes in sheep [73]. Further analysis from Chip-seq and RNA-Seq data will be required to uncover the function and regulatory mechanisms of *PCDH9* to adapt to hot and humid environments in sheep.

## Conclusions

In summary, we conducted a comprehensive and integrated analysis of Asiatic mouflon and domestic sheep based on WGS, mitogenome and Y-chromosomal genetic variants. Our results suggested that the mitochondrial haplogroups C/E and Y-chromosomal haplogroup HY2 of domestic sheep could have originated from the Asiatic mouflons in Iran by introgression and further clarified the effects of the Iranian Asiatic mouflon in the domestication processing. Additionally, our results revealed the introgression signals of the Asiatic mouflons in Iran in several domestic sheep populations in South and Southeast Asia and showed that the introgression could have facilitated their adaptation to hot and humid environments near the Indian Ocean. Our findings added new insights into the domestication and subsequent evolutionary history of sheep.

## Declarations

### Ethics approval and consent to participate

Not applicable.

### Consent for publication

Not applicable.

### Availability of data and materials

The whole genome re-sequence data used for the study is publicly available under the sample accession numbers listed in Additional file 1 Table S1. All mitogenomes assembled in this study were deposited in NCBI’s SRA under the accession number OR160429-OR161034. All scripts used for this work were performed using open-source software tools and are available from the corresponding authors upon request.

## Authors’ contributions

FHL and MHL conceived and supervised the study. DFW and FHL conducted the data analysis. DFW, FHL and POW wrote the manuscript, and MHL and POW revised the paper. All authors read and approved the final manuscript.

## Acknowledgements

We thank the High-performance Computing Platform of China Agricultural University for providing computing resources.

## Additional files

### Additional file 1 Table S1

Format: excel

Title: 780 samples information for autochromosome analysis.

### Additional file 1 Table S2

Format: excel

Title: 770 samples information for mitochondria chromosome analysis.

### Additional file 1 Table S3

Format: excel

Title: 337 samples information of *Cyt-b* sequence in this study.

### Additional file 1 Table S4

Format: excel

Title: 239 samples for Y chromosome analysis in this study.

### Additional file 1 Table S5

Format: excel

Title: Kinship of 32 *O. gmelini* samples.

### Additional file 1 Table S6

Format: excel

Title: The *F*ST of autosome, mitochondria and Y chromosome of *O. gmelini*_1, *O. gmelini*_2 and *O. aries*.

### Additional file 1 Table S7

Format: excel

Title: The nucleotide diversity / haplotype diversity of autosome, mitochondria and Y chromosome of *O. gmelini*_1, *O. gmelini*_2 and *O. aries*.

### Additional file 1 Table S8

Format: excel

Title: Haplotype information of mitogenomes and Y chromosomes.

### Additional file 1 Table S9

Format: excel

Title: Split times estimated by SMC++.

### Additional file 1 Table S10

Format: excel

Title: Haplotype information of *Cyt-b* sequences.

### Additional file 1 Table S11

Format: excel

Title: *D*-statistics from Iranian mouflon to SSA populations.

### Additional file 1 Table S12

Format: excel

Title: Details for chromosome-level introgression across four breeds.

### Additional file 1 Table S13

Format: excel

Title: Common introgressed blocks from *O.gmelini*_2 into four SSA populations.

### Additional file 1 Table S14

Format: excel

Title: Population branch statistic (PBS) of putative selective regions in the SSA population.

### Additional file 1 Table S15

Format: excel

Title: Overlaps regions among selected regions identified by PBS, introgression regions and QTLs regions.

### Additional file 2 Figure S1

Format: word

Title: 780 autosome NJ-tree showing the same two cluster of *O. gmelini* as Fig.1b and domestic sheep samples clustered based on their geographic distribution.

### Additional file 2 Figure S2

Format: word

Title: Admixture of 770 samples for *K*=2-9.

### Additional file 2 Figure S3

Format: word

Title: Admixture of SSA 18 breeds with two *O. gmelini* populations for *K*= 2-9.

